# Tigecycline pharmacodynamics in the hollow fiber system of *Mycobacterium avium*-complex lung disease, and the utility of MICs and time-kill studies in drug development

**DOI:** 10.1101/2025.07.29.667481

**Authors:** Devyani Deshpande, Shashikant Srivastava, Tawanda Gumbo

## Abstract

**Background:** Guideline-based therapy (GBT) drugs for *Mycobacterium avium-complex* (MAC) lung disease (LD) were chosen in part because they have low MICs. Despite these low MICs, GBT achieves six-month sustained sputum culture conversion in only 43% of patients.

**Methods:** First, we co-incubated tigecycline with MAC for seven days in time-kill studies and calculated the exposure mediating 50% of maximal effect (E_max_) or EC_50_. Next, we performed tigecycline exposure-effect studies in the hollow fiber system of MAC (HFS-MAC) inoculated with the reference ATCC#700898 isolate. Third, we performed an exposure-effect study in the HFS-MAC inoculated with 5 different isolates. Finally, the target exposure (EC_80_) was used to identify a clinical dose of inhaled tigecycline for MAC-LD in 10,000 subject Monte Carlo experiments (MCE).

**Results:** In time-kill studies the EC_50_ was 0-24h area under the concentration-time curve-to-MIC (AUC_0-24_/MIC) of 62.24 for extracellular and 0.14 for intracellular MAC (p<0.001). In the HFS-MAC inoculated with ATCC#700898, the EC_50_ statistically differed between sampling days by 2,370.7%. However, studies with five different isolates demonstrated a stable and robust day-to-day EC_50_ (%CV=18.18%), with an EC_80_ AUC_0-24_/MIC of 33.65. The E_max_ was 4.84 log_10_ CFU/mL. In MCE, tigecycline inhalational doses of 35-40 mg/day achieved the EC_80_ target in >90% of virtual patients, with and an MIC breakpoint of 256 mg/L.

**Conclusion:** Time-kill studies do not inform on PK/PD target exposures or extent of kill. Inclusion of multiple MAC isolates in HFS-MAC studies improves precision of pharmacokinetic/pharmacodynamic parameter estimates. Tigecycline via the inhalational route could contribute to treatment of MAC-LD.

## 1. Introduction

Guideline-based treatment (GBT) of *Mycobacterium avium complex* (MAC) lung disease (LD) of a macrolide plus rifamycin plus ethambutol achieves six-month sustained sputum culture conversion rates (SSCC) in only 43-53% of patients (Kwak et al., 2017; Pasipanodya et al., 2017; Daley et al., 2020). Refractory MAC-LD is defined as failure to achieve SSCC after six months of GBT, which means most patients fail the GBT (Daley et al., 2020). The best predictors of poor SSCC and mortality on treatment with GBT are a high MAC bacterial burden at the start of therapy (*B_0_*) and the presence of cavities >2 cm in diameter (Lam et al., 2006; Kang et al., 2021; Magombedze et al., 2021). The effect of *B_0_* on treatment outcomes is consistent with the inoculum effect in pharmacokinetics (PK) and pharmacodynamics (PD) science. For cavity size, drug concentrations and penetration are known to fall proportional to the diameter, which means centers of large cavities have poor drug concentrations (Lam et al., 2006; Dheda et al., 2018; Kang et al., 2021; Magombedze et al., 2021). The intracellular hollow fiber system model of MAC-LD (HFS-MAC) can mimic the *B_0_* in patient cavities, the intracellular nature of MAC in patients, and human-like antibiotic intrapulmonary PKs to mirror the SSCC reported in patients on GBT (Deshpande et al., 2010b; Schmalstieg et al., 2012; Deshpande et al., 2016; Deshpande et al., 2017a; Deshpande et al., 2017c; Ruth et al., 2019; Deshpande et al., 2020; Deshpande et al., 2023; Deshpande et al., 2024a; Deshpande et al., 2024b; Singh et al., 2024). Previously, GBT in the HFS-MAC killed between 0 and 2.3 log_10_ CFU/mL below *B_0_*, but kill below *B_0_* was observed in only 40% of the clinical isolates tested (Deshpande et al., 2023). Here, we used the HFS-MAC to test tigecycline for potential use in MAC-LD.

The first screening test for candidate drug activity for MAC is MICs. Based on the definition of resistance as an MIC ≥8 μg/mL, MICs for MAC clinical isolates for three tetracyclines (minocycline, tigecycline, omadacycline) led to the conclusion that 100% of *M. avium* and >87% of *M. intracellulare* isolates were resistant (Li et al., 2023). In a study by Wallace et al, 100% of 11 isolates were resistant to tigecycline and minocycline (Wallace et al., 2002). In our collection of 44 MAC isolates, resistance to tigecycline was in 90.9%, to omadacycline in 88.6%, and to minocycline in 86.4% of all isolates (Singh et al., 2025). Therefore, the general perception is that tetracyclines are “inactive” or have “no *in vitro* activity” against MAC, and thus would be futile to test these drugs for MAC-LD (Wallace et al., 2002; Li et al., 2023). However, despite these high MICs, minocycline and omadacycline monotherapies demonstrated better microbial kill than GBT in the HFS-MAC (Ruth et al., 2019; Brown-Elliott and Wallace, 2021; 2022; Chapagain et al., 2022; Li et al., 2023). Therefore, here we tested tigecycline in the HFS-MAC.

## 2. Methods

### 2.1. Materials and isolates

For MICs, static concentrations exposure-effect studies in 12-well tissue culture plates, and first HFS-MAC we used *M. avium* (American Type Culture Collection 700898, ATCC#700898).

Another five clinical MAC strains (two *M avium*, and three *M intracellulare*) were used in validation HFS-MAC studies. These five different MAC isolates, reflect the same response rates in the HFS-MAC as seen in patients on treatment with GBT (Deshpande et al., 2023). THP-1 monocytes (ATCC TIB-202) were used in 12-well studies. Phorbol myristate acetate (PMA), RPMI-1640 medium, and heat-inactivated fetal bovine serum (FBS), were purchased from Sigma-Aldrich (St. Louis, MO). Tigecycline was purchased from Baylor University Medical Center, Pharmacy (Dallas, TX, USA). Cellulosic HFS cartridges were purchased from FiberCell (Frederick, MD, USA).

### 2.2. MICs

We utilized the standard broth microdilution method using cation-adjusted Mueller-Hinton broth (CAMHB) supplemented with 5% oleic acid-albumin-dextrose-catalase (OADC) to determine the MICs (CLSI, 2018). The MIC assays were as described by the Clinical Laboratory Standards Institute, and our recent publications (CLSI, 2018; Singh et al., 2025). MICs were tested over a concentration range of 0.06 mg/L to 64 mg/L.

### 2.3. Static “time-kill” concentration-effect against extracellular and intracellular MAC

ATCC#700898 cultures in log phase growth at a bacterial density of ∼10^5^ CFU/mL were co-incubated with tigecycline at concentrations of 0 to 128mg/L in Middlebrook 7H9 broth supplemented with 10% OADC. On day 7, samples were washed, serially diluted, and inoculated onto Middlebrook 7H10 agar supplemented with 10% OADC. CFUs were recorded after incubation at 37°C for 10 days. The experiment was performed with three replicates per concentration.

For intracellular studies, THP-1 monocytes were cultured and infected with ATCC#700898 as described previously (Deshpande et al., 2010b; Deshpande et al., 2016; Chapagain et al., 2022; Deshpande et al., 2023). Adhered and infected cells were co-incubated with tigecycline dissolved in RPMI at final concentrations of 0-8 mg/L for 7 days. THP-1 cells were then lysed, as described before, followed by quantitation on Middlebrook 7H10 agar (Deshpande et al., 2010b; Deshpande et al., 2016; Chapagain et al., 2022; Deshpande et al., 2023). The experiment was performed with three replicates per concentration.

### 2.4. Hollow fiber system model of MAC

#### 2.4.1. Exposure-effect with ATCC#700898

HFS-MAC units were set up with a *B_0_* mimicking the bacterial burden in cavitary disease in 20mL peripheral compartment, as described in the past (Deshpande et al., 2010a; Deshpande et al., 2010b; Schmalstieg et al., 2012; Deshpande et al., 2016; Deshpande et al., 2017a; Deshpande et al., 2017c; Ruth et al., 2019; Deshpande et al., 2020; Chapagain et al., 2022; Deshpande et al., 2023; Deshpande et al., 2024a; Deshpande et al., 2024b; Singh et al., 2024). We mimicked the intra-pulmonary PKs of tigecycline based on a median epithelial lining fluid (ELF) to-plasma ratio of 1.5-2.4, and an ELF and alveolar macrophages half-life of 24-39h, respectively, and alveolar macrophage-to-ELF AUC ratio of 58.77 (Conte et al., 2005; De Pascale et al., 2014; Gotfried et al., 2017; Dimopoulos et al., 2022). Tigecycline doses were prepared fresh each day. Seven tigecycline doses were administered once daily with target AUCs of 0 to 480 mg*h/L with 2-fold decrease in doses. Nontreated HFS-MAC units served as growth controls. The target tigecycline half-life (t_1/2_) was 30h, midway between 24-39h reported in literature. The HFS-MAC units were repetitively sampled on days 2, 7, 14, 21, and 26 for THP-1 count and bacterial burden estimation. Bacterial burden was determined by lysing THP-1 cells, as described above. Samples were inoculated on Middlebrook 7H10 agar and in the MGIT tubes to record time-to-positivity (TTP) using the Epicenter Software. The central compartments were sampled on the last day of the study at pre-dose (0h) and 1, 6,12, and 24h post dosing. Tigecycline concentrations were measured using the method described previously (Ferro et al., 2016; Deshpande et al., 2019).

#### 2.4.2. Exposure-effect with five clinical MAC isolates

For drug development purposes, the US Food and Drug Administration (FDA) and European Medicines Authorities (EMA) require testing at least 4-5 clinical isolates for robust PK/PD target setting (EMA, 2016). Therefore, we performed an exposure-effect HFS-MAC study with the five different clinical MAC isolates. The five isolates we used were chosen based on our interactions with regulatory authorities who wanted a panel of 5 isolates that reflected the heterogeneity in response to GBT encountered in patients, to allow generalizability of findings. Each isolate had its own non-treated control (AUC/MIC=0) and received a single daily dose of one of 4 different AUCs; given the different MICs this led to a total of six AUC/MIC ratios among the 5 isolates (including the AUC/MIC=0). Sampling for bacterial burden and PKs were as in section 2.4.1.

### 2.5. PK/PD modeling and

We used the inhibitory sigmoid maximal effect (E_max_) model on each sampling day for bacterial burden (either TTP or log_10_ CFU/mL) versus drug exposures (concentration or AUC/MIC). The four parameters for this model are E_max_, the effective concentration mediating 50% of E_max_ or EC_50_, the Hill slope of H, and bacterial burden in non-treated controls (E_con_). From this, we calculated the EC_80_ as target exposure for MCEs. Further details on the MCEs are described in online supplementary methods.

### 2.6. Monte Carlo experiments (MCEs)

Tigecycline has been formulated for both intravenous and inhalational formulation, the latter by three separate groups in search for less systemic toxicity (Petersen et al., 1999; Himstedt et al., 2022; Nair and Smyth, 2023). For intravenous tigecycline dosing we used Gotfried et al (Gotfried et al., 2017). For inhaled tigecycline we used the epithelial lining fluid (ELF) concentrations and systemic pharmacokinetic parameters generated for humans in the semi-mechanistic physiology-based pharmacokinetic model derived by Himstedt et al (Himstedt et al., 2022). Unlike Himstedt et al, we assumed zero % of drug dose would go down the gastrointestinal tract and be absorbed that way, but that all absorption would be in respiratory tract. For inhaled therapy, the ELF compartment was specified as the central compartment, the systemic circulation peripheral compartment 1, and the rest of the tissues as peripheral compartment 2. The best compartmental model was then chosen based on Akaike Information Criteria. The model was then used for MCE.

The inter-individual variability (as %CV) in Himstedt et al in the inbred rats was 4.5% for clearance and 5.36% for volume, whereas in the study by Gotfried et al in people were 17.75% for clearance and 21.26% for volume; we set these at 25% in our MCE (Gotfried et al., 2017). We examined both the intravenous route and the inhalational therapy route for doses of 0, 1, 10, 20, 30, 40, 50, 100 and 200 mg once daily. Pharmacokinetic parameters and covariance matrix were added to subroutine PRIOR in ADAPT 5, and the ELF and plasma concentration-time profiles generated for each dose in 10,000 virtual subjects; the values of these estimates are shown side by side in the results section. The AUCs so generated were examined at each MIC for probability target attainment (PTA) to achieve EC_80_ is ELF (Singh et al., 2025). Regarding toxicity, AUCs at each dose were examined for a target plasma AUC_0-24_ of >6.87 mg*h/L, demonstrated by Rubino et al to be associated with a higher probability of nausea and vomiting (Rubino et al., 2012).

## 3. Results

### 3.1. MICs and static tigecycline concentration versus effect

The tigecycline MICs for ATCC#700898, and the five clinical isolates were as shown in **Table 1**. Trailing effect was observed for all isolates. One isolate had an MIC of 8mg/L, considered resistant.

**Table 1.** Tigecycline Minimum Inhibitory Concentrations.

| **Isolate** | **Minimum inhibitory concentration (mg/L)** | **Trailing effect observed over** |
| --- | --- | --- |
| **ATCC#700898** | 2 | 3 tube-dilutions |
| **S1** | 1 | 3 tube-dilutions |
| **S2** | 2 | 3 tube-dilutions |
| **S3** | 1 | 3 tube-dilutions |
| **S4** | 8 | 3 tube-dilutions |
| **D5** | 4 | 3 tube-dilutions |

**Tabel 2** shows the tigecycline inhibitory sigmoid E_max_ model parameter estimates and microbial kill below day bacterial burden (*B_0_*), for the extracellular and intracellular ATCC#700898 time-kill studies. Notably, the microbial kill below *B_0_*, H, and the EC_50_ differed significantly between intracellular versus extracellular assays, with the EC_50_ differing by a factor of 531 between the two conditions.

**Table 2.** Inhibitory Sigmoid E_max_ Parameters for intracellular versus extracellular MAC in time-kill studies.

|  | **Extracellular** | **Intracellular** |
| --- | --- | --- |
| **$E_{\text{con}}$ $\log_{10}$ CFU/mL** | 10.83±0.23 | 7.19±0.12 |
| **$E_{\text{max}}$ $\log_{10}$ CFU/mL** | 10.0±0.37 | 4.33±0.1 |
| **Kill below $B_0$ at $E_{\text{max}}$** | 1.45±0.05 | 4.46±0.17 |
| **H** | 1.38±0.14 | 0.38±0.17 |
| **EC<sub>50</sub> mg/L** | 5.31±0.43 | 0.01±0.02 |
| **EC<sub>80</sub> mg/L** | 14.5±1.17 | 0.38±0.77 |
| **$r^2$** | 0.99 | 0.98 |

### 3.2. HFS-MAC exposure-effect results with MAC ATCC#700898

The tigecycline concentrations measured in the central compartment of each HFS-MAC unit are shown in **Figure 1A**, as are the AUC_0-24_/MIC achieved for each regimen. PK modeling revealed an elimination rate constant (k_el_) of 0.023±0.003h^-1^, a volume of 364±33 mL, and a t_1/2_ of 29.9±3.7h in the HFS-MAC. Thus, the %CV of the final PK parameters between HFS-MAC replicates was ∼10%. The observed versus model-predicted concentrations are shown in **Figure 1B**. The measured tigecycline concentrations versus PK model predicted concentrations had a slope of 0.95 (95% CI:0.92-0.99) for a one compartment model (r^2^=0.98) indicating minimal bias. These PK results were used to calculate the AUC_0-24_/MIC achieved for each regimen.

**Figure 1.**
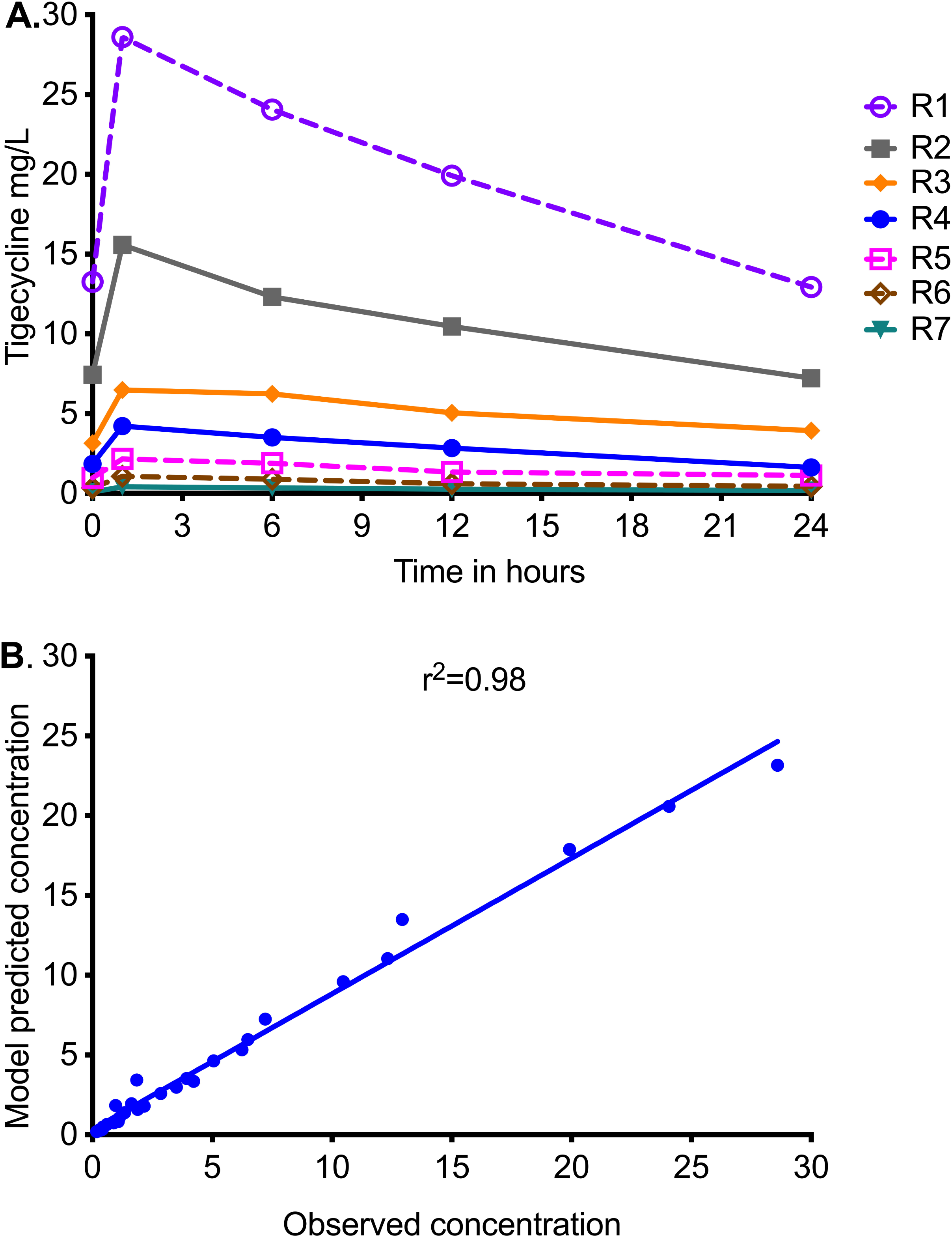
Tigecycline concentration-time profiles achieved in HFS-MAC units. R1 to R7 are descending doses of tigecycline, administered once a day. **A.** Symbols are concentrations achieved at the different time points in each HFS-MAC. The line graphs shown are point-to-point of the observed concentrations. **B.** The measured tigecycline concentrations versus PK model predicted concentrations.

**Figure 2A** shows the sampling day-to-day bacterial burden changes using the TTP readout from the MGIT. TTP goes up as bacterial burden decreases, and for the non-treated TTP decreased throughout the 26 days of study. All exposures demonstrated a biphasic effect except in the highest dose (AUC/MIC=240.8), an initial kill as demonstrated by increasing TTP, followed by rebound growth shown by decreasing TTP. The time-to-positivity readout was able to separate out the exposure-response of the four highest exposures from each other at time points before day 14. **Figure 2B** shows results using CFU/mL readout. The *B_0_* was 6.8 log_10_ CFU/mL. All exposures, except non-treated controls or AUC/MIC=0, killed below *B_0_* and then rebounded. The highest exposure killed 4.9 log_10_ CFU/mL below *B_0_*. However, the CFU/mL readout was able to separate the exposure-response of the four lowest exposures from each other at time points before day 14.

**Figure 2.**
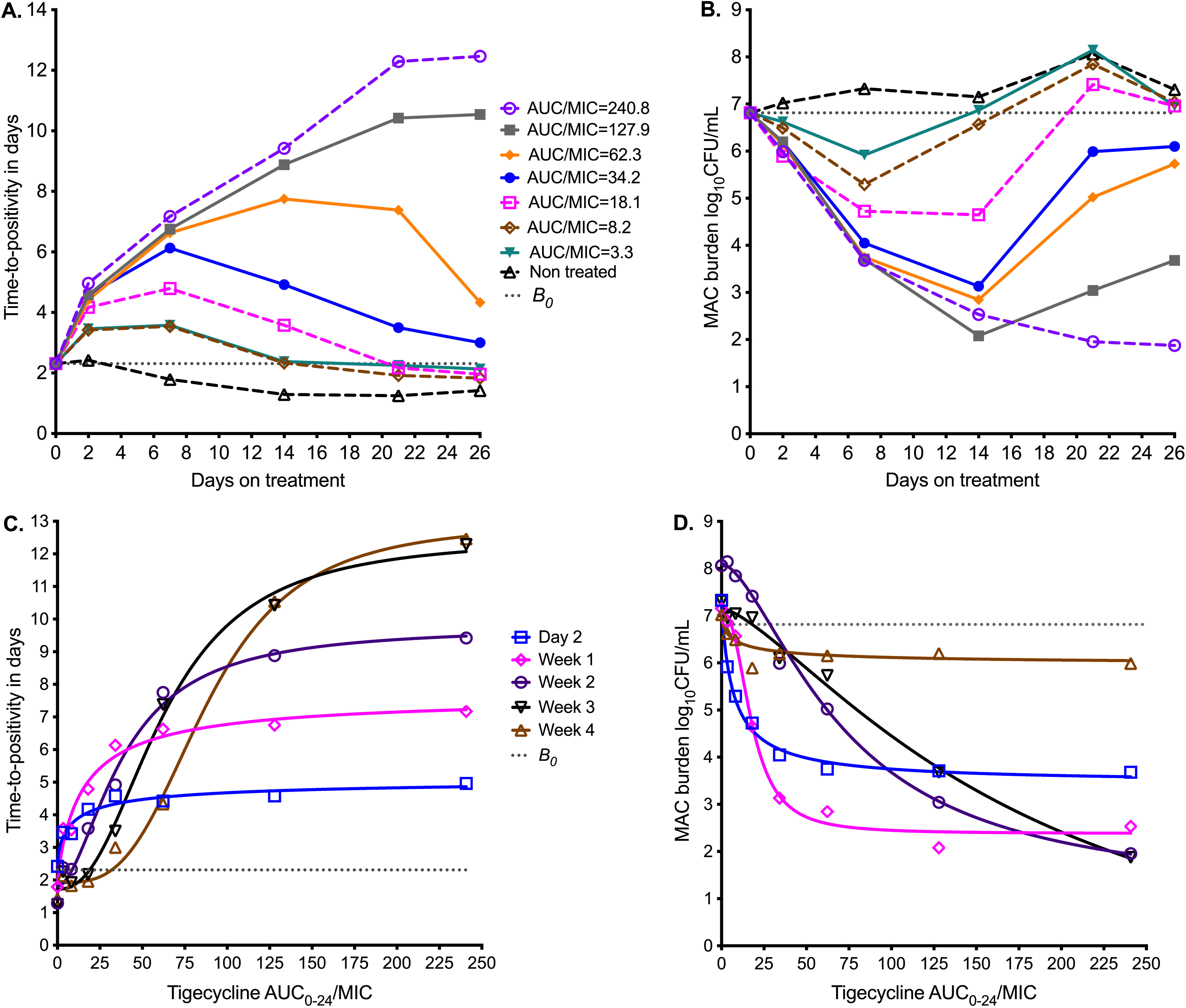
Tigecycline PK/PD at different sampling points in the HFS-MAC. Symbols are bacterial burden measurements observed at the different time points in each HFS-MAC **A.** Change in bacterial burden from sampling day-to-sampling day based on time-to-positivity readout, for each AUC/MIC exposure. The higher the time-to-positivity the lower the bacterial burden. **B** Colony forming units per mL (log_10_) readout demonstrate the same pattern. **C.** Inhibitory sigmoid E_max_ model using time-to-positivity as a pharmacodynamic parameter. **D**. Inhibitory sigmoid E_max_ model using log CFU/mL as pharmacodynamic parameter readout.

Inhibitory sigmoid E_max_ modeling results using TTP readout are shown in **Figure 2C** and those using CFU/mL in **Figure 2D**. The parameter estimates for both readouts are summarized in **Table 3**. **Table 3** shows that the EC_50_ and H changed multiple folds between sampling days and often differed by bacterial burden readout. For TTP readout, EC_50_ changed from an AUC_0-24_/MIC of 9.49 on day 7 to 88.4 on day 26 (931.5%-fold change); the H also changed in the same period. For CFU/mL readout, EC_50_ changed from an AUC_0-24_/MIC of 7.58 on day 7 to 179.70 on day 26 (2,370.7%-fold change). A simple naïve pooled average of all EC_50_ for all sampling days for both TTP and CFU/mL readouts was an AUC_0-24_/MIC of 49.90±54.4, which translates to a mean EC_80_ AUC_0-24_/MIC of 126.4 (95%CI: 30.45-222.3), and thus an imprecise estimate.

**Table 3.**
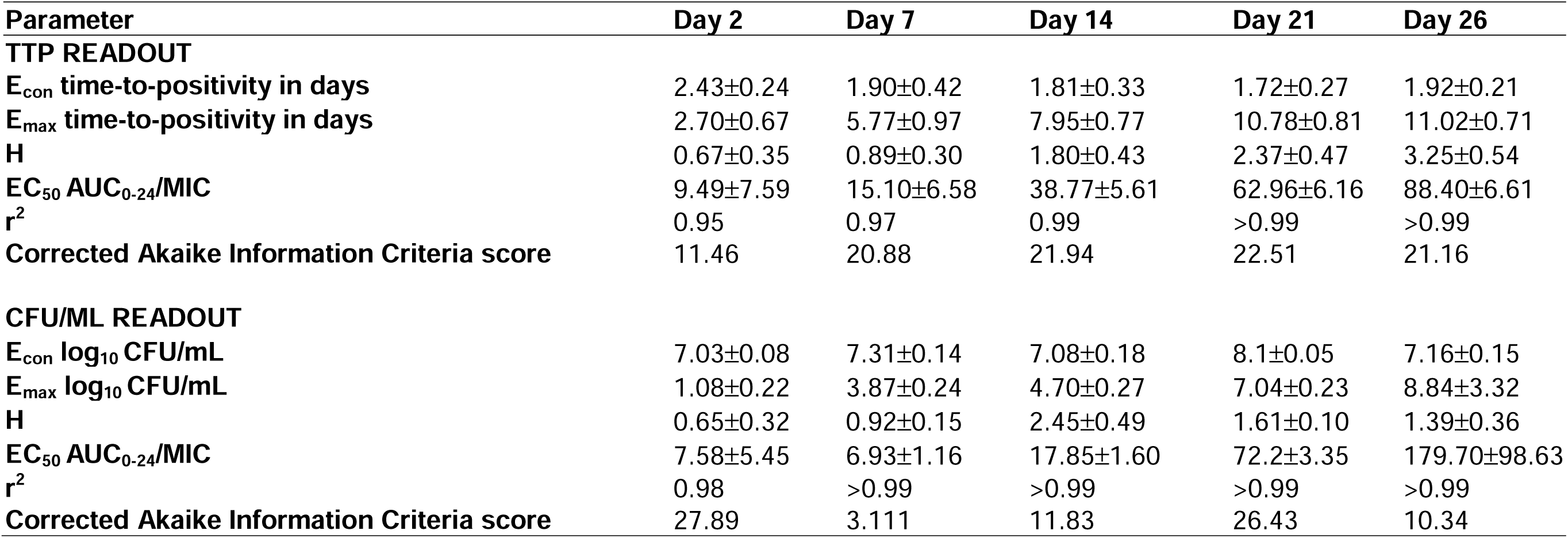
Pharmacokinetic/pharmacodynamic parameter estimates and standard error using two readouts.

### 3.3. HFS-MAC exposure-effect results with five clinical strains

**Figure 3A** shows the results of PK modeling for the HFS-MAC units with five clinical strains, from just before first dose till steady state. The MICs of each isolate are shown in **Table 1**.

**Figure 3.**
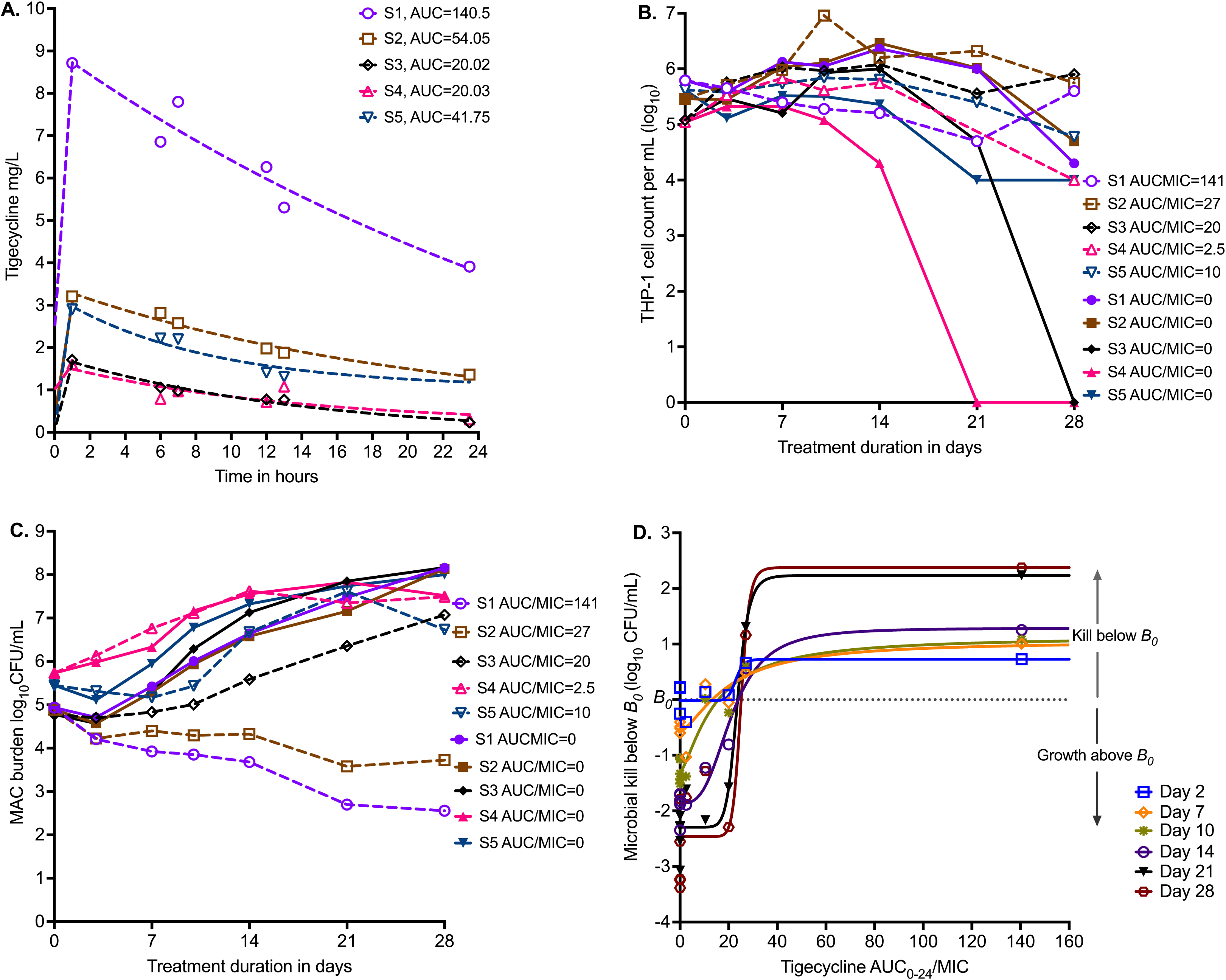
Tigecycline PK/PD in the HFS-MAC with 5 MAC strains. **A.** Tigecycline 24h concentration-time profiles based on measurement in each HFS-MAC unit. The line graphs are pharmacokinetic model based output. Non-treated controls all had 0 concentration and are not shown. **B.** Infected THP-1 cells are killed by MAC, based on the virulence of the isolate. Non-treated systems, that is AUC/MIC=0, are shown as solid symbols **C.** Infectivity is reflected by variability in *B_0_*. Effect of each of five different exposures (one for each isolate) versus no drug treatment on day-to-day changes in bacterial burden is shown. **D.** No single isolate was treated with a full dose-response range; they all received a single daily AUC/MIC exposure (in addition to each having a non-treated controls). Shown is inhibitory sigmoid E_max_ model that used data from all five isolates pooled for each sampling day (i.e., co-modeled), with pharmacodynamic readout of kill below *B_0_*.

**Figure 3B** shows the viability of THP-1 cells during the experiment, with the non-treated HFS-MAC illustrating the heterogeneity of the different MAC isolates in killing the THP-1 cells. **Figure 3C** shows that the *B_0_* ranged from 4.8 to 5.73 log_10_ CFU/mL. Thus, despite the same multiplicity of infection, the *B_0_* differed between the isolates because of differences in infectivity. The growth patterns in non-treated controls also differed by isolate. **Figure 3C** shows changes in bacterial burden for each clinical strain versus the tigecycline AUC_0-24_/MIC achieved in respective HFS-MAC units.

We co-modeled all 5 isolates (each one receiving a different AUC/MIC exposure) in one inhibitory sigmoid E_max_ equation. Given differences in *B_0_*and E_con_ (bacterial burden in non-treated controls) we used microbial kill below *B_0_* for each isolate for each sampling day, with results shown in **Figure 3D** and **Table 4**. **Table 4** demonstrates virtually no change in EC_50_ from sampling day-to-day with high precision (18.18 %CV). The average EC_80_ across sampling days was an AUC_0-24_/MIC of 33.65 (95% CI: 23.73-43.96), a much narrower confidence interval than in the ATCC#700898-inoculated HFS-MAC experiment.

**Table 4.** Inhibitory sigmoid E_max_ estimates with kill below B_0_ as pharmacodynamic parameter.

| | $E_{\text{con}} \log_{10} \text{CFU/mL}$ | | $E_{\text{max}} \log_{10} \text{CFU/mL}$ | | H | | $EC_{50} \text{AUC}_{0-24}/\text{MIC}$ | | $r^2$ | AICc |
| --- | --- | --- | --- | --- | --- | --- | --- | --- | --- | --- |
|  | Estimate | SEM | Estimate | SEM | Estimate | SEM | Estimate | SEM |  |  |
| **Day 2** | -0.02 | 0.12 | 0.74 | 0.22 | 14.18 | 20.10 | 22.91 | 4.95 | 0.75 | 0.48 |
| **Day 7** | -0.57 | 0.12 | 1.62 | 0.33 | 1.50 | 1.18 | 19.89 | 8.50 | 0.87 | -10.31 |
| **Day 10** | -1.31 | 0.13 | 2.45 | 0.31 | 1.39 | 0.65 | 14.24 | 4.22 | 0.94 | -10.00 |
| **Day 14** | -1.86 | 0.12 | 3.15 | 0.25 | 3.12 | 1.21 | 21.29 | 2.26 | 0.96 | -9.60 |
| **Day 21** | -2.29 | 0.16 | 4.53 | 0.33 | 10.08 | 3.10 | 23.63 | 1.16 | 0.97 | -2.49 |
| **Day 28** | -2.46 | 0.30 | 4.84 | 0.64 | 14.70 | 17.39 | 25.09 | 2.57 | 0.91 | 11.91 |

SEM- standard error of the mean

### 3.4. Monte Carlo Experiments (MCE) of intravenous and inhalation tigecycline doses

The PK parameter estimates and variance in domain of input are compared to MCE generated values in virtual subjects in **Table 5** and demonstrate good recapitulation of parameters reported in patients. We examined tigecycline doses of 1, 10, 20, 30, 40, 50, 100, and 200 mg once daily for ability to achieve or exceed EC_80_ in ELF. The probability of target attainment (PTA) for the intravenous doses up to 1, 10, 20, 30, 40, 50, 100 and 200 mg were zero within the tigecycline MIC range examined, which was based on the MIC distribution from our work elsewhere (Singh et al., 2025).The PTAs for inhaled doses were as shown in **Figure 4A**. The inhalation doses of 1 mg and 10 mg showed that PTA falls below 90% before the modal MIC of 128 mg/L, while the dose of 40 mg achieved a PTA of >90% at that MIC. The clinical susceptibility breakpoint for the dose of 30 mg/day is an MIC of 128 mg/L, and for 40 and 50 mg/day an MIC of 256 mg/L.

**Figure 4.**
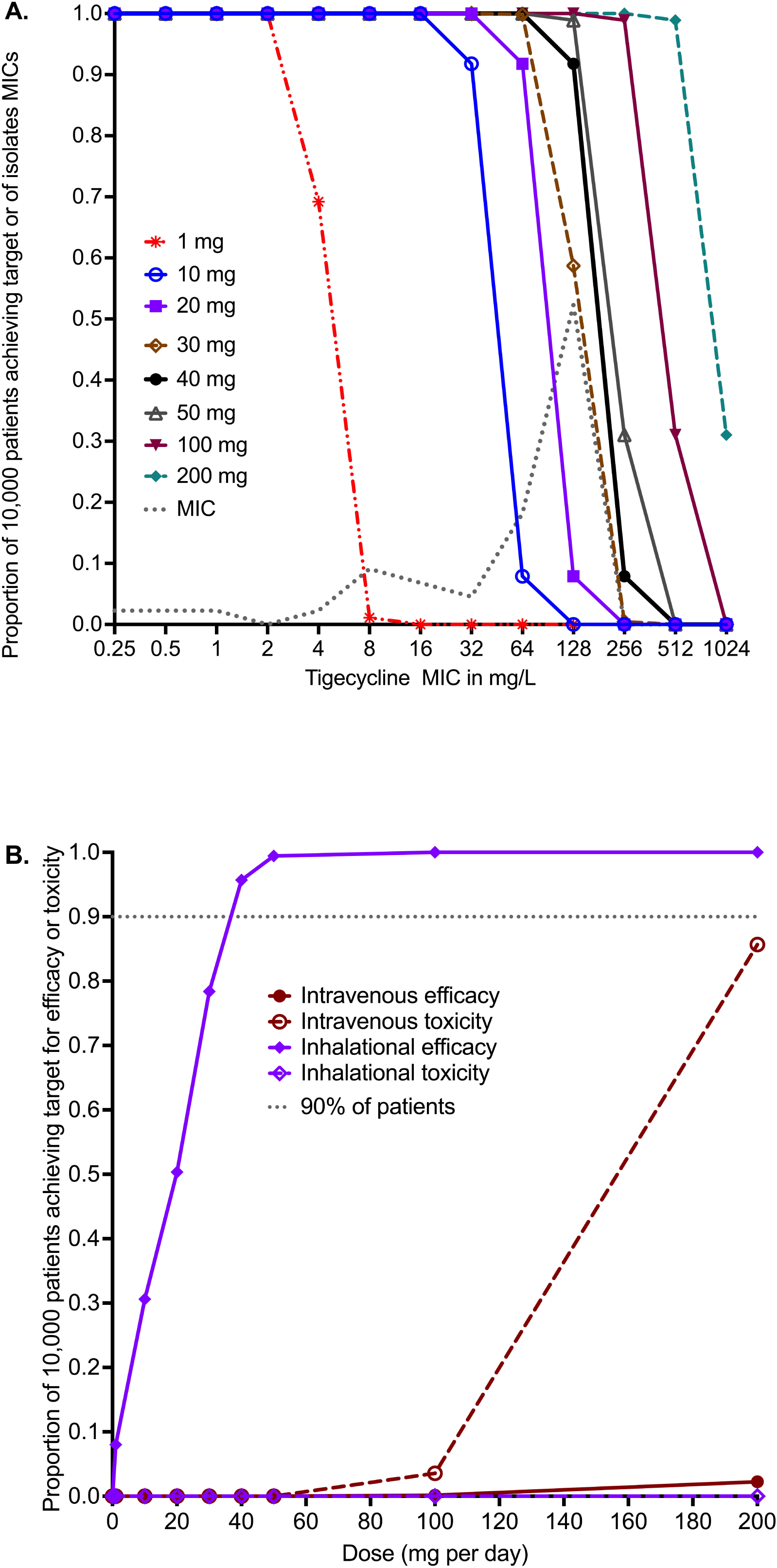
*in silico* dose finding for intravenous and inhaled tigecycline. **A.** Probability of target attainment (PTA) for inhaled tigecycline doses. **B.** The cumulative fraction of response (CFR) for intravenous dosing versus inhaled dosing for both efficacy and toxicity thresholds.

**Table 5.** Monte Carlo simulation experiment pharmacokinetic model output versus domain of input values.

|  | Parameters in domain of input |  | Output in 10,000-subject simulation |  |
| --- | --- | --- | --- | --- |
|  | Parameter estimate | IIV (%) | Parameter estimate | IIV (%) |
| **ELF Clearance in L/h** | 0.68 x10<sup>-2</sup> | 25.0 | 0.68 x10<sup>-2</sup> | 25.24 |
| **ELF Volume in L** | 0.07 x10<sup>-2</sup> | 25.0 | 0.07 x10<sup>-2</sup> | 24.98 |
| **K<sub>a</sub> (from lung to systemic)** | 1.002 | 25.0 | 0.99 | 39.45 |
| **Systemic clearance L/h** | 23.1 | 25.0 | 22.97 | 25.10 |
| **Systemic Volume in L** | 315 | 25.0 | 314.6 | 25.31 |
| **Lung AUC<sub>0-24</sub> for inhaled 10 mg dose (mg\*h/L)** |  |  | 1,520 | 25.19 |
| **Lung AUC<sub>0-24</sub> for inhaled 30 mg dose (mg\*h/L)** |  |  | 4,559 | 25.19 |
| **Lung AUC<sub>0-24</sub> for inhaled 40 mg dose (mg\*h/L)** |  |  | 6,079 | 25.19 |

IIV=inter individual variability

The cumulative fraction of responses (CFR) for the lung AUC_0-24_/MIC≥33.55 are shown in **Figure 4B** for both intravenous and inhalational doses. The intravenous dose of 200 mg/day achieved the EC_80_ target in ELF in only 2.26% of 10,000 virtual patients, lower than the 1 mg/day inhalational doses which achieved the CFR in 8.49% of patients. **Figure 4B** shows that 90% proportion of patients would achieve the target at a dose of 35 mg/day, and 95.70% at 40 mg/day. No inhalational doses tested in the MCE was predicted to achieve the plasma AUC_0-24_ of >6.87 mg*h/L associated with a higher probability of toxicity. The intravenous dose of 100 mg/day was predicted to achieve the AUC associated with higher risk of adverse effects in 3.57% of patients, whereas the toxicity probability with 200 mg/day dose increased to 85.67% in the 10,000 virtual patients.

## 4. Discussion

Like other bacteria, the first step in MAC drug development program is testing for MICs. Based on that, and based on susceptibility breakpoint of 8 mg/L, >90% of MAC clinical isolates were considered resistant to tigecycline, suggesting that the drug has poor activity and should not be used for MAC-LD (Wallace et al., 2002; Li et al., 2023; Singh et al., 2025). Such susceptibility studies were also used to make decisions for use of minocycline and omadacycline, which were considered resistant. However, despite these high MICs, in an open-label clinical trial of minocycline combined with clarithromycin and clofazimine, SSCC rate was 64%, which is better than the GBT (Roussel and Igual, 1998). Moreover, both minocycline and omadacycline monotherapies demonstrated better microbial kill than GBT in the HFS-MAC (Ruth et al., 2019; Brown-Elliott and Wallace, 2021; 2022; Chapagain et al., 2022; Li et al., 2023). The discrepancy between high MICs and good efficacy against MAC by minocycline, tigecycline, and omadacycline is likely due, in part, to two “technical” reasons. First, the chemical instability or degradation rate of tetracyclines in growth medium is faster than MAC doubling time (Chapagain et al., 2022; Singh et al., 2025). Second, the discrepancy could be because MICs are performed using extracellular assays, while in patients’ lungs MAC is predominantly intracellular (Ushiki et al., 2011; Hibiya et al., 2012). Minocycline, tigecycline and omadacycline achieve alveolar cell-to-serum AUC penetration ratios of 25-77-fold (Conte et al., 2005; Gotfried et al., 2017). In our static concentration experiments, the tigecycline EC_50_ for intracellular MAC was 531-fold lower than for extracellular MAC, suggesting that extracellular assays may not be as predictive as regards to potency against MAC.

The findings of MIC distribution-based “resistance” not predicting PK/PD response seem to extend to other antibiotic classes. There are high MICs for MAC but good response to ceftriaxone in the intracellular HFS-MAC, while for clarithromycin and azithromycin good MICs with >90% of isolates in the “susceptible” range are what has given us the current poor response rates to GBT (Schmalstieg et al., 2012; Kwak et al., 2017; Pasipanodya et al., 2017; Daley et al., 2020; Deshpande et al., 2023). Thus, conventional MICs, which use extracellular MAC, could underestimate drug potency for intracellular pathogens. This means that MIC distribution range could be misleading as a decision point as to whether a drug will work in the treatment of MAC-LD or not. Given the poor efficacy of GBT drugs in patients and in the HFS-MAC despite “good” MICs, this begs the question: *have we been throwing away the babies and keeping the bath water* (Schmalstieg et al., 2012; Kwak et al., 2017; Pasipanodya et al., 2017; Deshpande et al., 2023)? Have we been ignoring some drugs in the tetracycline class because of high MICs while keeping the current GBT drugs with “good” MICs but demonstrably poor efficacy in the clinic?

As part of a more complete PK/PD data package, guidance by regulatory authorities includes co-incubation of bacteria with static concentrations of the candidate drug, including time-kill curves (EMA, 2016). For MAC we have proposed use of intracellular and extracellular “time-kill” assays, as a more “PK/PD” approach than MIC (Deshpande et al., 2017b). From a PK/PD standpoint, the question is if such parameter values as EC_50_, or EC_80_, identified in “time-kill” assays contribute to those identified in the HFS-MAC (or mice for that matter) or in patients. As an example, in **Table 2**, if the concentrations are converted to AUC_0-24_/MIC the EC_80_ for extracellular becomes 174 and that for intracellular 4.56. Both figures differ from the EC_80_ of AUC_0-24_/MIC of 50.12 on day 7 in the HFS-MAC. This is important for two reasons. First, the only directly transferable information between the preclinical models and patients are the PK/PD targets (EC_50_ and EC_80_). Second, the whole point of PK/PD studies is identification of target exposure and dosing schedule for clinical application. It is not possible to perform dose-fractionation using these static concentration assays. Therefore, we question the PK/PD utility of performing these static concentration assays in the drug development process for MAC.

Here, as elsewhere, we found that the EC_80_ target values with ATCC#700898 isolate “wobble” and vary across a wide range between sampling days (Musuka et al., 2013; Chapagain et al., 2022; Deshpande et al., 2023). This “wobble” makes it difficult to identify the target exposure value to use in dose selection. Our results here and elsewhere, and from others, suggest that the ATCC#700898 isolate likely provides a misestimate of PK/PD target exposure. Similarly, in the HFS-MAC inoculated with ATCC#700898, GBT combination killed 2.1 log_10_ CFU/mL below *B_0_* but the effect on five clinical isolates was such that GBT killed below *B_0_*in only 2/5 strains and failed in the remaining three, very similar to what is encountered in patients (Kwak et al., 2017; Pasipanodya et al., 2017; Daley et al., 2020; Deshpande et al., 2023). Here, tigecycline killed 4.9 log_10_ CFU/mL ATCC#700898 below *B_0_*, versus less effect in the five clinical isolates in the HFS-MAC. This means that similar to rapidly growing bacteria, and both US FDA and EMA Guidance, HFS-MAC studies should include more than four isolates for a more precise estimation of PK/PD target exposures (EMA, 2016).

Finally, using the EC_80_ target from the five clinical strains in the HFS-MAC study, we identified intravenous and inhalational tigecycline doses for the treatment of MAC-LD. The intravenous doses performed poorly. An inhaled dose of 35-40 mg/day was able to achieve the exposure target in >90% of the 10,000 virtual patients, with predicted fewer gastrointestinal side effects. This is where the MIC distribution should be used, to identify optimal dose and PK/PD-based breakpoints, and not as an arbiter for which new drug should be developed for the treatment of MAC. The MIC distribution (by extension MIC_90_) may be high, but if a drug is administered in such a way that high enough concentrations can be achieved in the lung, then the “high” MIC is not a reason to reject a candidate drug for PK/PD investigations for the treatment of MAC-LD.

There are several limitations in our studies. First, given the degradation of tigecycline in agar, we could not characterize the PK/PD parameters associated with resistance suppression. Second, we did not employ a full dosing schedule to enact a complete exposure-response surface for each of the five clinical isolates. However, we borrowed the technique used in murine studies by Craig and Andes, for rapidly growing Gram-negative bailli and Gram-positive cocci (Craig and Andes, 2008). They first identified a full exposure response using the standard laboratory strain, then used a limited number of doses in a larger number of clinical isolates with different MICs and then co-model all results in a single inhibitory sigmoid E_max_ model. Third, in our MCE we used ELF PKs based on a PBPK model, and not from direct sampling of ELF in patients after administration of inhalational doses. However, even when we used the clearance rates based on rat ELF PKs from direct sampling or observations (usually faster clearance than achieved in people), the PTAs did not change significantly.

## 5. Conclusion

Here, we examined tigecycline potency and efficacy using MICs in the literature, static concentrations versus exposure, and in the intracellular HFS-MAC. First, MIC distributions which showed >90% of isolates would be resistant, and static concentration versus effect studies, were not informative as to tigecycline microbial kill below *B_0_* and PK/PD target exposures in the HFS-MAC. Second, an inhaled tigecycline of 35-40 mg/day achieved target exposures in lungs in MCEs and should be tested as therapy for MAC-LD, after further preclinical studies. Third, we found a PK/PD susceptibility breakpoint MIC of 256 mg/L with inhaled dosing. This means the range of tigecycline MIC determination should be extended to at least two dilutions above 256 mg/L, for inhaled dosing. Finally, the reference ATCC#700898 gave an over-optimistic and imprecise PK/PD target exposure, when compared with HFS-MAC inoculated with at least five MAC isolates. Thus, multiple MAC strains are required for a precise PK/PD target estimation.

## Acknowledgement.

None.

## Conflict Of Interest

Nothing to declare.

## Funding Source

Tawanda Gumbo received funding for this study from the National Institute of Allergy and Infectious Diseases of the National Institutes of Health (R56 AI111985). Shashikant Srivastava is supported by 1R21AI148096 and 1R01AI179827 grants from the National Institute of Allergy and Infectious Diseases (NIAID), KANT23G0 from the Cystic Fibrosis Foundation, and NTM Education and Research funding support from the University of Texas at Tyler.

## Data Availability Statement

The raw data for the results presented in the manuscript is available with the corresponding authors upon a reasonable request.

## Ethical Approval

Not applicable.

## Author Contributions

Conceptualization and design: DD and TG. MIC determination experiments: DD. HFS-MAC experiments: DD, SS, TG. PK/PD modeling, mathematical simulations: TG. TG wrote the first draft of the manuscript. All authors reviewed and approved the final version of the manuscript.

## Use of AI to write manuscript

The authors did not use any AI during the preparation of this work.

## Notes

### Competing Interest Statement

The authors have declared no competing interest.

